# Better antibodies engineered with a GLIMPSE of human data

**DOI:** 10.1101/2025.06.08.658113

**Authors:** N. Lance Hepler, Andrew J. Hill, David B. Jaffe, Michael C. Gibbons, Katherine A. Pfeiffer, Denise M. Hilton, Melanie Freeman, Wyatt J. McDonnell.

## Abstract

GLIMPSE-1 is a protein language model trained solely on paired human antibody sequences. It captures immunological features and achieves best-in-class performance in humanization benchmarks. We demonstrate the utility of GLIMPSE-1 in humanization; engineering of antibodies for affinity, species cross-reactivity, and key developability parameters; and the creation of highly divergent functional variants with <90% sequence identity to a marketed antibody. Learning exclusively from human antibody data enables GLIMPSE-1 to enhance therapeutics and native antibodies based on patterns in the human repertoire.

**Disclaimer:** While we provide detailed descriptions of experimental methods and success metrics, certain methodological details of GLIMPSE-1 remain proprietary and/or redacted in this work for commercial considerations. We warmly invite researchers and potential collaborators interested in accessing GLIMPSE-1 to connect with our team via.

## 1 Introduction

Therapeutic antibodies are a cornerstone of modern medicine, with applications in many diseases. Their success stems from their natural function as products of the adaptive immune system, shaped by evolution for millions of years to recognize and respond to targets with extraordinary selectivity. This specificity, combined with favorable stability and safety characteristics, underpins their widespread therapeutic success.

Therapeutic antibody discovery traditionally relies on biological systems, primarily through model organism immunization and display libraries from immunized animals. Newer approaches, including our own Anthrobody^®^ platform, leverage human biology to discover antibodies with superior properties *de novo*. Human antibodies contain unique sequence patterns and structural motifs adapted to human physiology, ensuring proper folding, stability, and immune compatibility. However, all discovery methods typically produce antibodies that require additional engineering to create sufficiently potent and manufacturable molecules. This need fuels ongoing advances in computational methods for antibody evaluation, engineering, and optimization.

We present GLIMPSE-1 (<u>G</u>enerative <u>L</u>anguage <u>I</u>mmunoglobulin <u>M</u>odel for <u>P</u>rotein <u>S</u>equence <u>E</u>ngineering), a protein language model trained exclusively on human antibodies to maintain the delicate balance achieved through evolution. This contrasts with prior work (**Table S1**) where models were commonly trained using heterogeneous datasets (PDB^1^, OAS^2^, and/or SAbDab^3^). Moreover, we use only paired sequences, to account for the biological and physical interactions between heavy chains and light chains. Notably, BALM-Burbach^4^ was also exclusively trained on paired human sequences, but data from only 4 donors were included. In further contrast, GLIMPSE-1 employs a different architecture and was trained on data from a much larger number of donors.

Here we show that GLIMPSE-1 mirrors human biology, enabling it to enhance existing therapeutics and antibodies found in humans. Our work bridges the gap between directly sourcing human antibodies and engineering them based on patterns observed in the broader human repertoire.

## 2 Results

### 2.1 GLIMPSE learns exclusively from paired human antibody data

We first trained and evaluated GLIMPSE-0, a RoBERTa^5^ model, using a corpus of unique, productive, and natively paired antibody Fv amino acid sequences, including public data from Jaffe *et al*. (2022) and proprietary data from Infinimmune’s Complete Human^®^ immunosequencing technology. GLIMPSE-0 produced surprisingly promising results despite being trained on data with limited diversity and suboptimal train/test splits. These early findings motivated us to develop a more robust model with an improved architecture and more diverse training data. Subsequently, we trained GLIMPSE-1, on a curated dataset also comprising public sequences from Jaffe *et al*. (2022) and proprietary sequences from Infinimmune’s Anthrobody^®^ and Complete Human^®^ platform technologies.

### 2.2 GLIMPSE-1 learns immunologically relevant sequence features of antibodies

Antibody Fv sequences develop through the rearrangement of variable (V), diversity (D), and joining (J) genes during B cell maturation. This process, along with somatic hypermutation (SHM), generates the sequence diversity that powers adaptive immunity. While this diversity is most pronounced at V(D)J junctions, most Fv sequences remain recognizably related to their original V gene sequences. A well-trained antibody language model (AbLM) should naturally recognize V gene lineages by clustering related superfamily and subfamily sequences while distinguishing between different ones. GLIMPSE-1 successfully captures this biological organization. When we embedded 0.6M paired Fv sequences from antibodies discovered at Infinimmune (see **Methods**), the resulting visualization revealed clear immunological structure (**Figure 1**).

**Figure 1:**
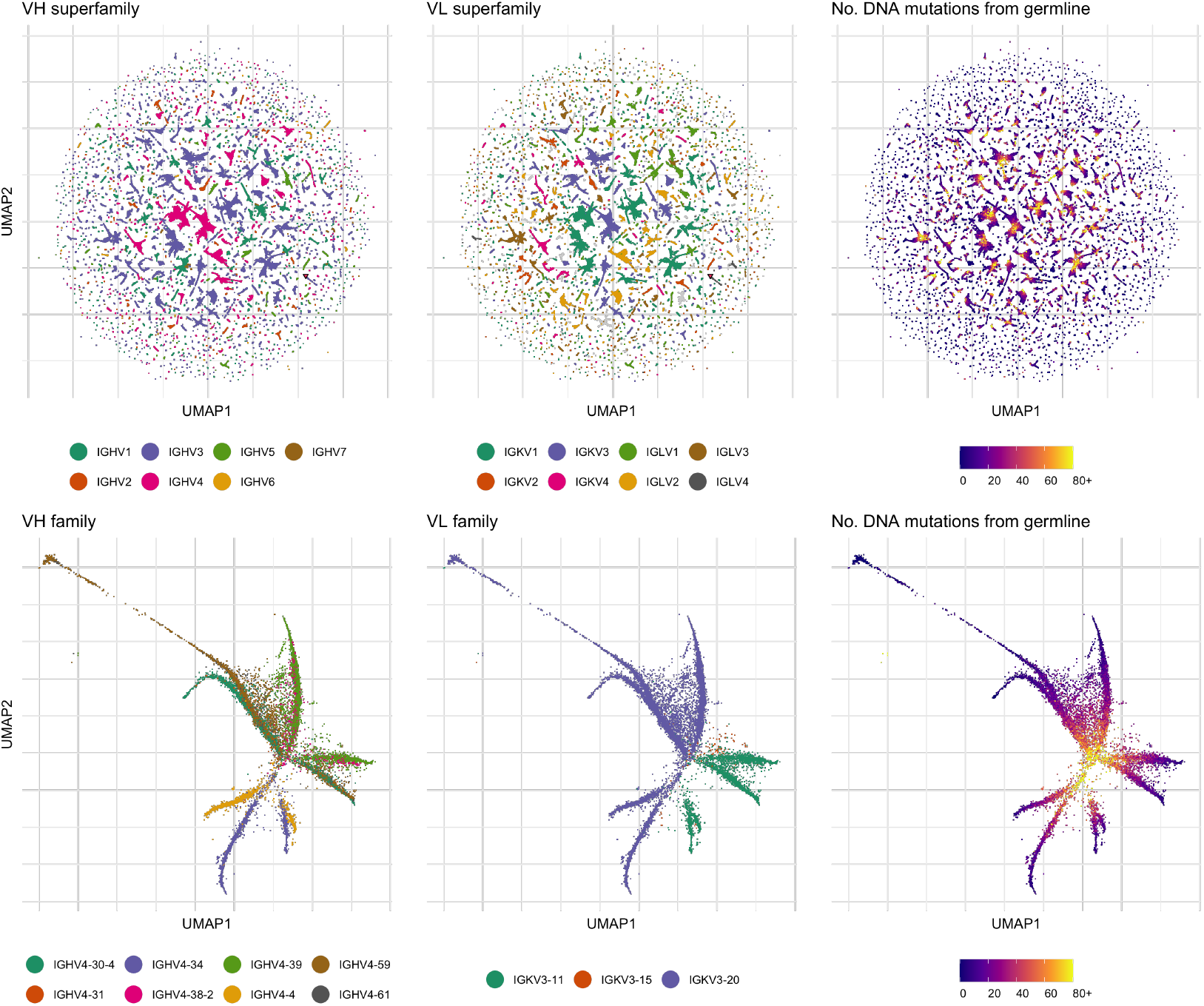
GLIMPSE-1 learns immunologically relevant sequence features of antibodies. Top: UMAP^7^ embeddings of GLIMPSE-1 last-layer embeddings for 0.6M antibodies discovered at Infinimmune colored by heavy chain V gene superfamily (left), light chain V gene superfamily (center), and number of DNA mutations from the inferred V gene allele for the donor of each particular antibody (right). Bottom: A single superfamily combination (IGHV4-IGKV3) from the global UMAP, showing how V gene subfamilies drive sequence clustering (left and center), while mutation accumulation (right) reduces subfamily identity while preserving superfamily association. This organization resembles a tree structure where identifiable subfamily sequences form distinct “branches” and more heavily mutated sequences gather in a central “trunk.”

The top row of **Figure 1** demonstrates this structure at the resolution of V gene superfamilies, with a curious pattern where more heavily mutated sequences appear more centrally within clusters. We examined this pattern more closely by focusing on the IGHV4 x IGKV3 superfamily cluster (bottom row). When colored by V gene subfamily, a tree-like structure emerges. Sequences clearly identifiable as specific subfamilies form distinct “branches”, while more heavily mutated sequences gather in a central “trunk” region having diverged further from their germline origins.

A biologically accurate AbLM should predict both which positions tolerate variation and which require conservation, beyond simply recognizing V genes. We analyzed GLIMPSE-1’s preferences across XY62, a TNFα-specific antibody isolated from a human donor using the Anthrobody^®^ platform. As shown in **Figure 2**, FR positions display strong conservation with GLIMPSE-1 favoring residues present in both germline and native antibody (purple). CDRs show greater diversity with complex preferences, sometimes favoring germline residues (blue), native non-germline residues (red), or alternative amino acids (grey). CDRH3 displays the highest diversity, consistent with its role in antigen specificity. These biases accurately reflect the biological constraints of antibody maturation.

**Figure 2:**
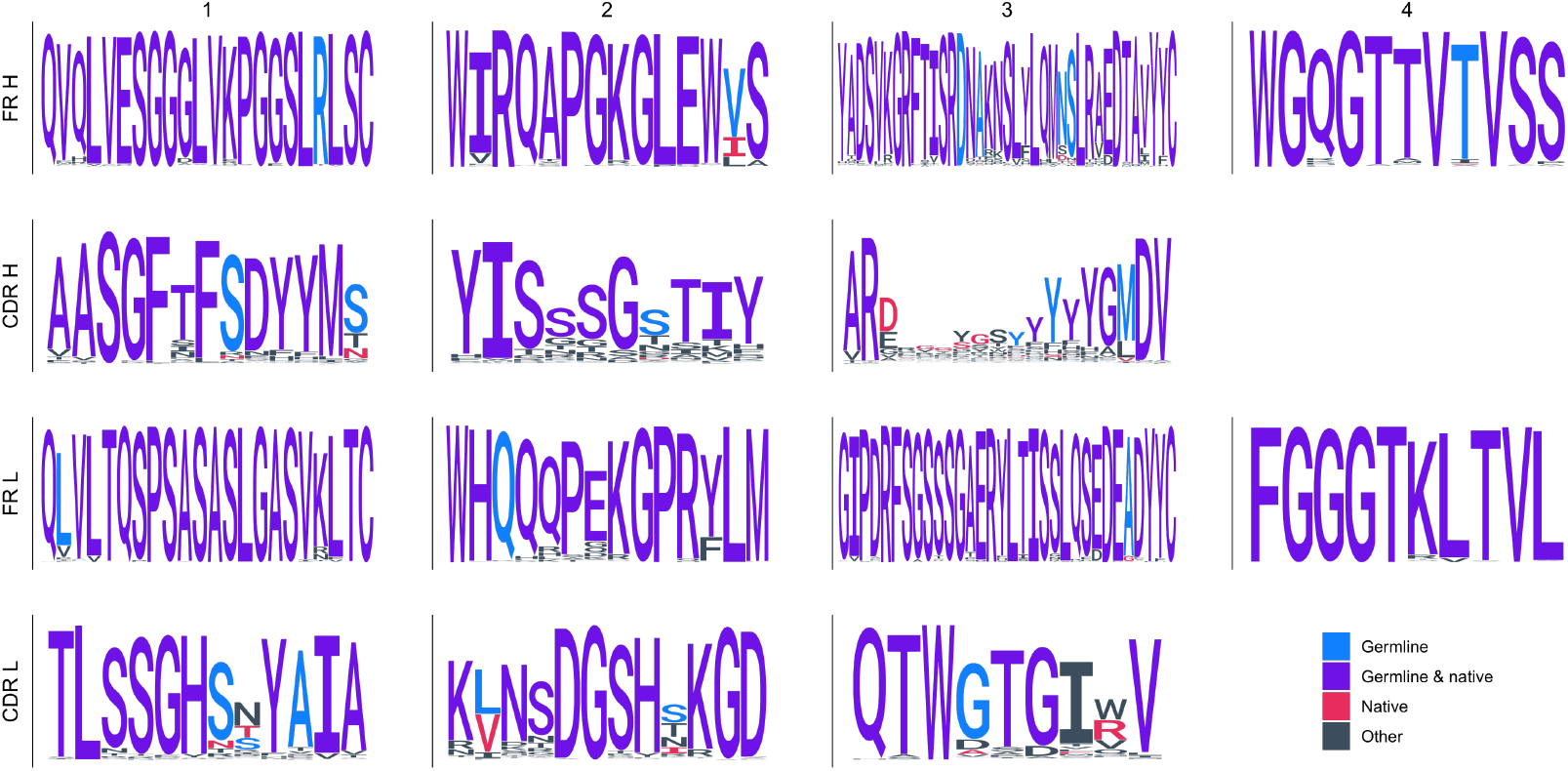
GLIMPSE-1 reflects freedoms and constraints of antibody sequence generation, maturation, and function. Sequence logos show GLIMPSE-1’s amino acid preferences across XY62, an anti-TNFα antibody. Colors indicate residue origin: germline non-native residues (blue), residues present in both native and germline sequences (purple), native non-germline residues (red), and alternative residues that are predicted by GLIMPSE-1 (grey). Stack height represents information content at each position, with taller stacks indicating stronger conservation. Framework regions (FR) show strong conservation with preference for germline-native residues, while complementarity-determining regions (CDRs) display greater sequence diversity and more complex preferences. CDRH3 shows the highest variability, consistent with its primary role in antigen recognition during antibody maturation.

### 2.3 GLIMPSE-1 achieves state-of-the-art performance in antibody humanization

Converting non-human or partially-human antibodies into effective therapeutics remains an important challenge in antibody development. Though modern humanization protocols have become more systematic, the process still requires careful engineering to preserve binding affinity while minimizing immunogenicity. Even with established methods, humanization projects require expertise and validation time, and outcomes can vary in how well they maintain the desired properties of the parent molecule while achieving human-like characteristics. AbLMs like GLIMPSE-1 may offer a new approach to potentially streamline this process while maintaining core antibody attributes.

We evaluated GLIMPSE-1’s humanization capabilities using 25 pairs of parental and humanized clinical molecules from Prihoda *et al*. [8], of which 20 could be independently confirmed. We benchmarked GLIMPSE-1 against nine other antibody language models using an iterative humanization approach (see **Methods**). As shown in **Figure 3**, GLIMPSE-1 matches or exceeds the performance of all other models, including Sapiens, in predicting the mutations found in clinical humanized antibodies. Notably, GLIMPSE-1 achieved this performance despite significantly more constrained training resources compared to Sapiens’, highlighting the value of a diverse and well-curated human dataset and its utility in training AbLMs.

**Figure 3:**
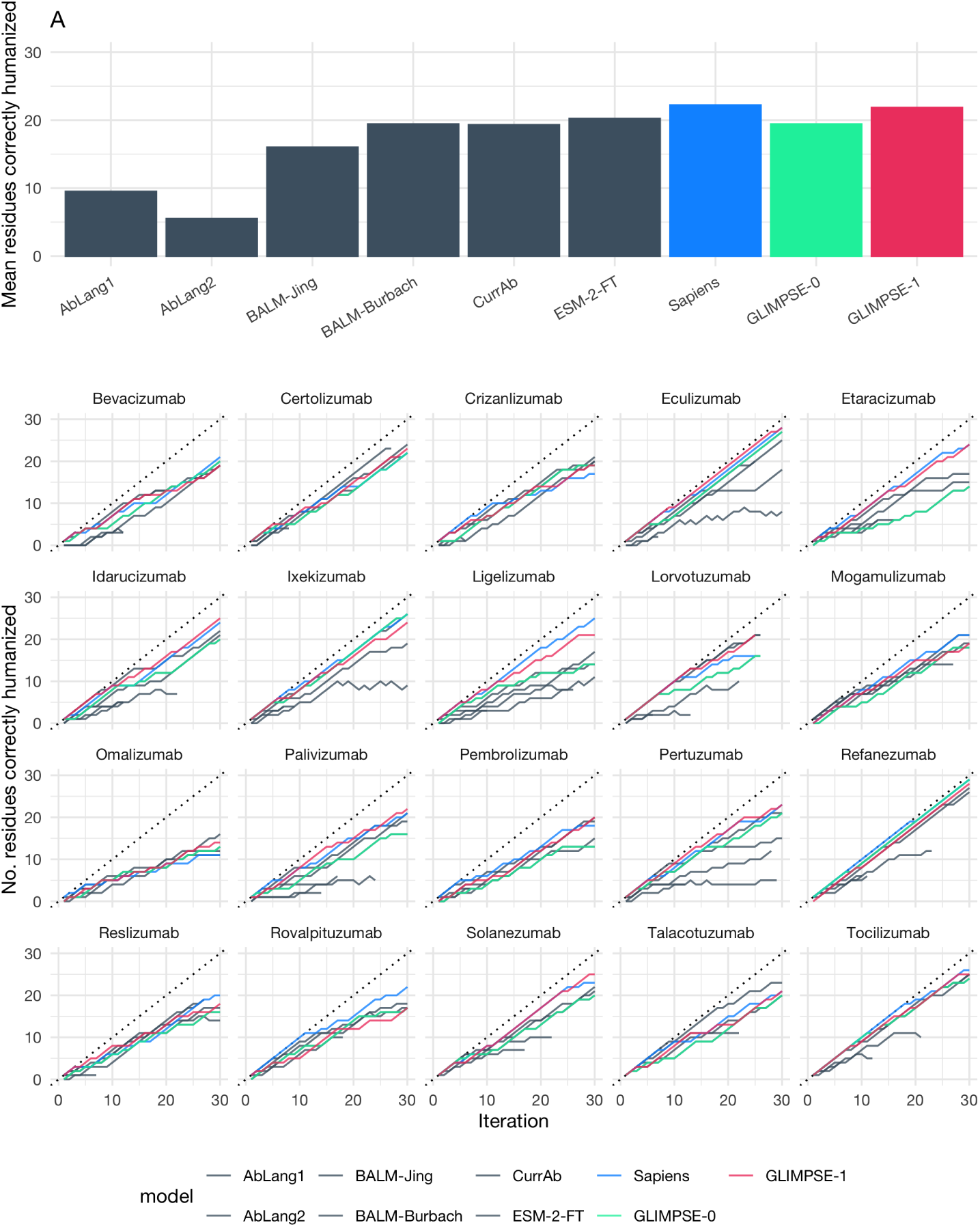
Humanization of 20 mouse-derived clinical stage antibody therapeutics. In each iteration, models were restricted to modifying only framework regions while preserving CDRs, selecting a single amino acid mutation aimed at humanizing the parental mouse antibody. (A) Mean number of residues correctly humanized by different antibody language models after 30 iterations. GLIMPSE-1 (red) and Sapiens (blue) are similarly capable of correctly predicting humanizing residues and outperform all other AbLMs evaluated. (B) Individual performance curves for each of the 20 antibodies. Each line represents a different model’s performance across iterations, with GLIMPSE-1 (red) and GLIMPSE-0 (green) consistently among the top performers. The dotted line represents a perfect 1:1 incorporation of mutations from parental mouse to humanized therapeutic sequence.

**Figure 3B** shows the performance of GLIMPSE-1 and other models across individual antibodies when we restricted models to modify only FRs while leaving CDRs intact. This approach mirrors common clinical practice to preserve antigen binding. We observed consistent strong performance from GLIMPSE-1 across the antibody panel, though the difficulty of replicating the original humanization strategies varied (cf. omalizumab vs. eculizumab). We obtained similar results when allowing modifications throughout the variable domains (**Figure S1**) or restricting changes to non-paratope residues (**Figure S2**).

### 2.4 GLIMPSE-1 exploits constrained diversity to engineer antibodies

We integrated GLIMPSE-1 into our antibody engineering process to optimize molecules against four undisclosed targets (A-D). We generated variant molecules using GLIMPSE-1 recommendations, complemented when available by intra-clonotype variation identified through our Complete Human^®^ technology.

SPR characterization revealed that while many variants showed reduced binding kinetics, several mutations for antibodies against each target significantly improved affinity compared to both parental molecules and biosimilars of clinical controls (**Figure 4**). Beyond affinity optimization, our engineering approach simultaneously addressed developability concerns by removing sequence liabilities predicted *in silico*, such as fragmentation, deamidation, and isomerization motifs, among others.

**Figure 4:**
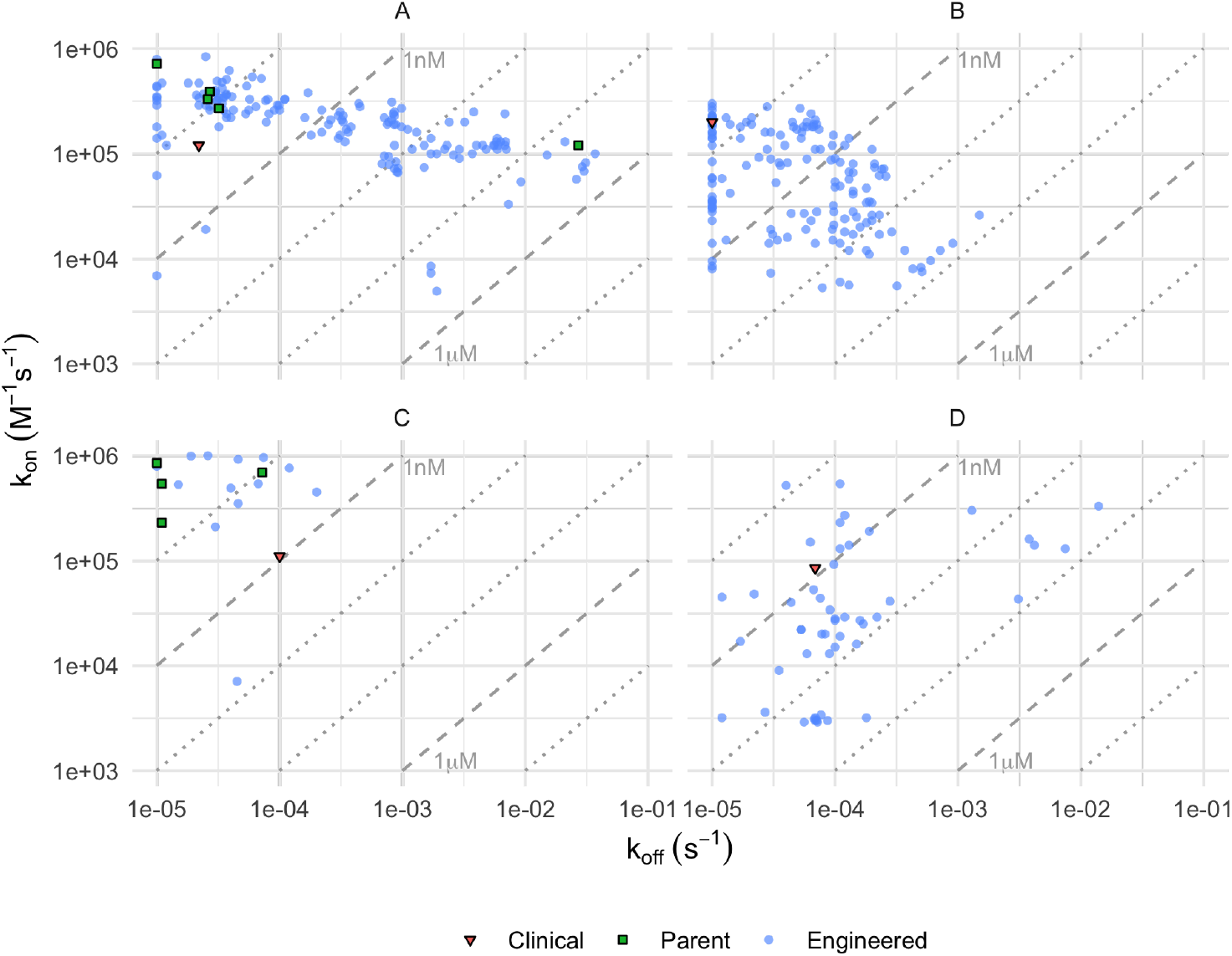
SPR affinity characterization of antibodies engineered using GLIMPSE-1. Kinetic parameters (k_off_ vs. k_on_) for antibodies targeting four undisclosed antigens (A-D). Colors indicate antibody categories: parent antibodies (green squares), engineered variants (blue circles), and clinical controls (red inverted triangles). Diagonal dashed lines mark constant K_D_ values (1nM and 1μM labeled). While many engineered variants exhibit reduced binding parameters, several variants demonstrate improved affinity compared to their respective parents and controls. We removed sequence liability motifs predicted *in silico* from these affinity-improved variants during the engineering process.

For Target A, we performed a deeper engineering campaign across five unique antibodies to mitigate potential liabilities while maintaining or improving binding. We evaluated these variants for binding to human, cynomolgus monkey, and mouse versions of the antigen (**Figure 5B, 5C**). As expected, binding affinity correlation was stronger between evolutionarily closer species (Cyno∼Human Pearson R^2^ = 0.78) than between more distant ones (Mouse∼Human Pearson R^2^ = 0.15). Variants appearing off-diagonal may represent cases where species-specific differences in the antigen epitope influence antibody recognition. Additionally, we engineered variants of families (INF1164, INF1169, and INF1181) for formulation-preferred isoelectric point (pI) as calculated using the method of Bjellqvist *et al*^9^. Binding affinity of these variants was minimally impacted by this engineering, as determined by bio-layer interferometry (**Figure 5D**).

**Figure 5:**
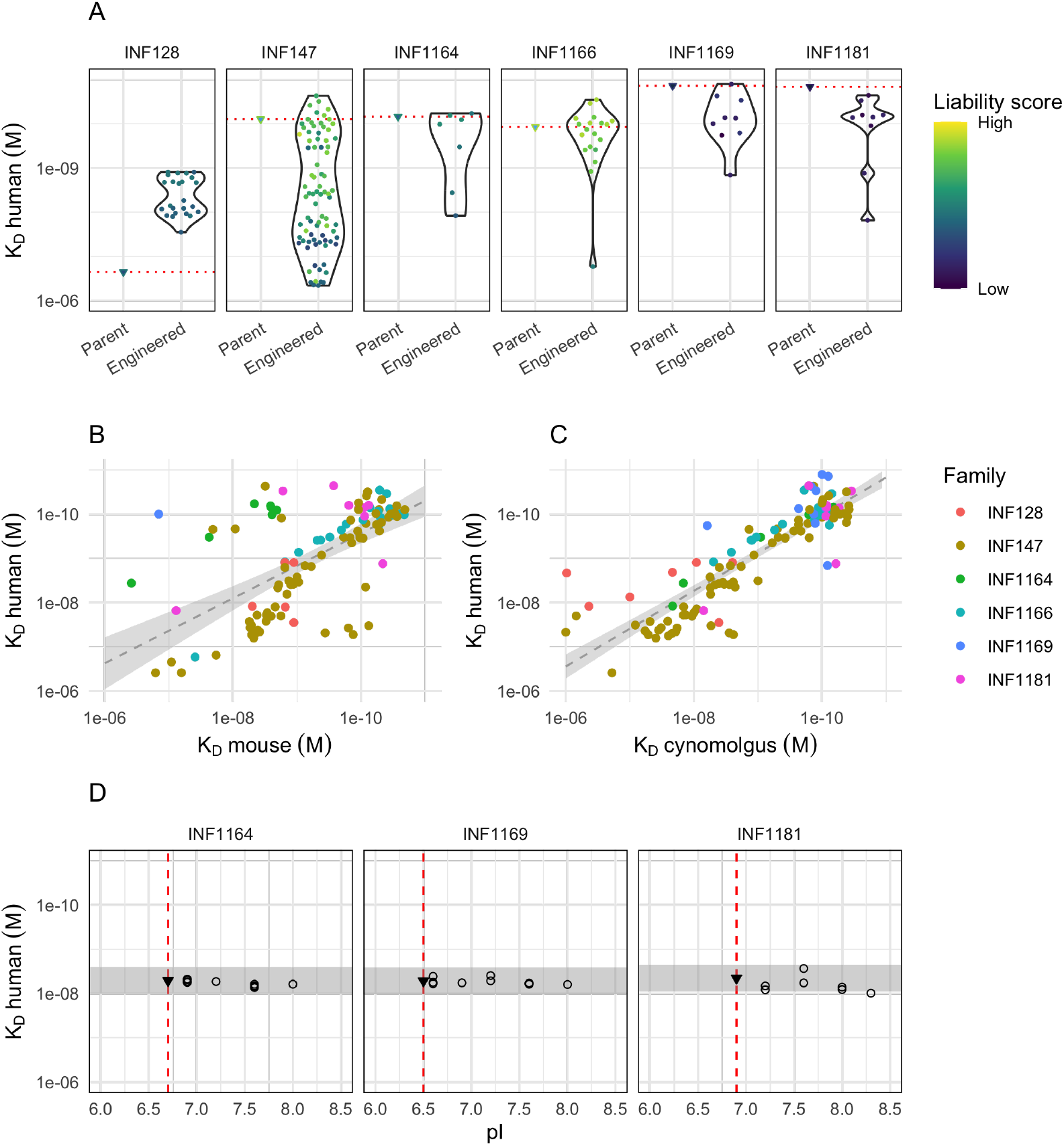
Sequence optimization of five unique antibodies that bind target A. (A) K_D_ values for parent antibodies (red dotted lines) and engineered variants (points) across five antibody families. Colors indicate liability scores from low (dark blue) to high (yellow). We mitigated high-risk sequence liabilities using GLIMPSE-1 predicted alternatives and clonally related sequences where available. (B-C) Correlation of binding affinity across species: human vs. mouse (B) and human vs. cynomolgus monkey (C). Points are colored by antibody family. The stronger correlation between human and cynomolgus (Pearson’s R^2^ = 0.78) compared to human and mouse (Pearson’s R^2^ = 0.15) reflects the evolutionary conservation of the antigen. Variants falling off the diagonal may represent antibodies binding to epitopes with species-specific differences. (D) Human binding affinity (K_D_) across predicted isoelectric point (pI) values for variants of antibodies INF1164, INF1169, and INF1181. Vertical dashed lines indicate the predicted pI value of the parental antibody, demonstrating successful engineering for formulation-preferred range. The grey band indicates a range of 2-fold relative to the K_D_ of the parental antibody.

**Figure 5A** reveals an important engineering principle: improving already high-affinity antibodies (e.g. INF1169, INF1181) presents a greater challenge than enhancing moderate-affinity starting points (e.g. INF147), demonstrating the diminishing returns often encountered in affinity maturation. While affinity optimization represents a common benchmark for antibody engineering platforms, GLIMPSE-1’s capacity to simultaneously address multiple design considerations points to its potential for solving more fundamental challenges in therapeutic antibody development.

### 2.5 GLIMPSE-1 streamlines identification of highly divergent and functional antibody variants

Therapeutic applications increasingly require antibody variants that preserve target binding while diverging significantly in sequence from the original molecule. Ideally these variants would occupy distinct viable sequence spaces while maintaining core functionality, akin to proteins that serve the same function across different species. Traditional methods to create such variants include CDR scanning mutagenesis, display libraries, epitope/paratope-guided design, and CDR grafting, each with limitations in speed and scope. In this setting, an AbLM could define the sequence constraints that permit substantial diversification from a parental antibody while maintaining essential functional attributes and desirable characteristics such as humanness.

GLIMPSE-1 offers a powerful approach for exploring novel functional sequence space starting from existing antibodies. We evaluated its capabilities using a clinically validated, marketed antibody that underwent humanization during its development (**Figure 6**). Our approach followed a two-stage engineering process.

**Figure 6:**
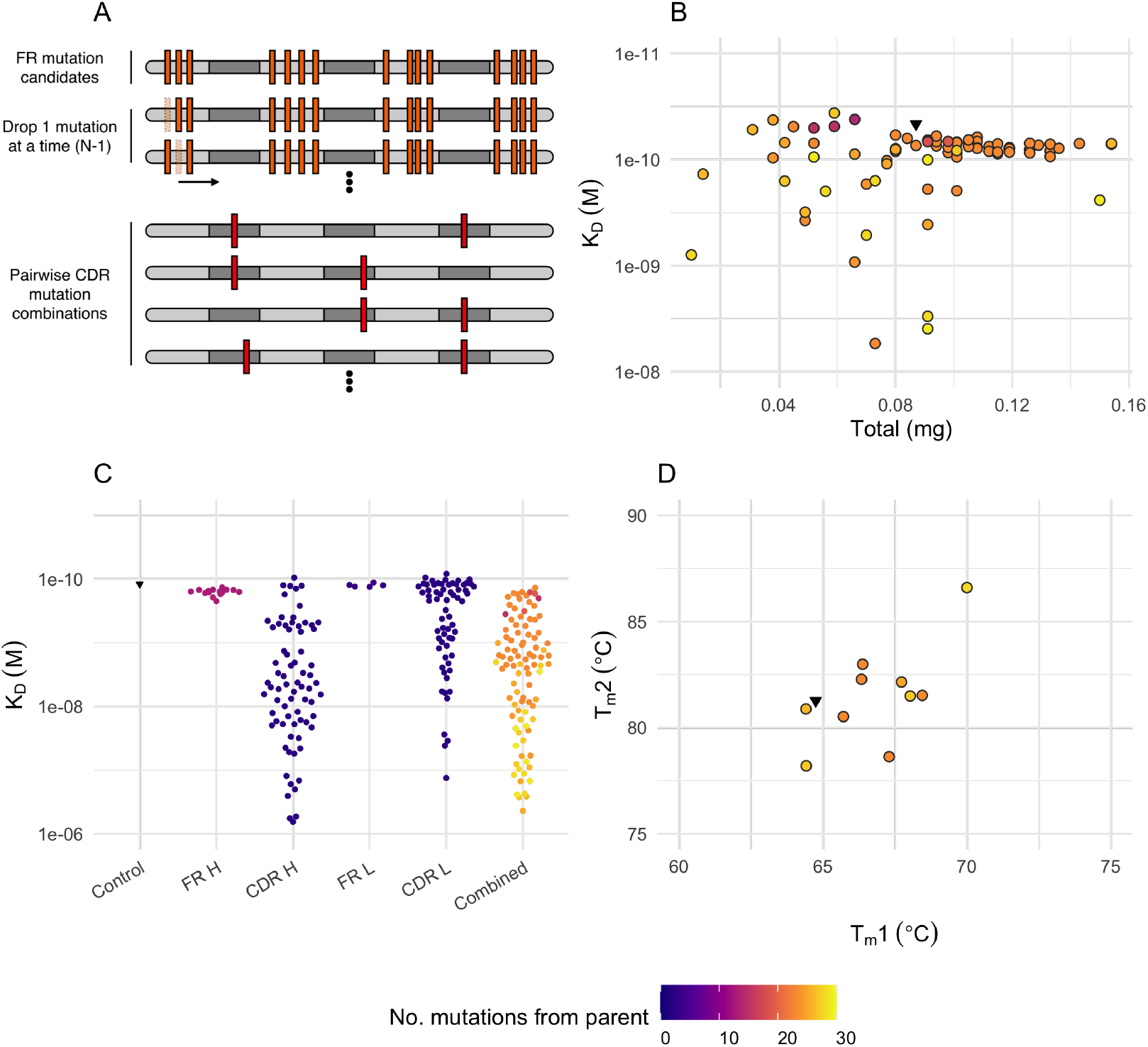
Engineering divergent, functional antibody variants. (A) Experimental design showing our two-stage approach in which we test multiple FR mutations per variant (top) and evaluate pairwise CDR combinations (bottom). (B) Expression yield vs. binding affinity for generated variants. (C) Binding affinity comparison across different mutation categories, with color indicating mutation count from parent sequence. (D) Thermal stability measurements (T_m_1 and T_m_2) showing preservation of stability in engineered variants.

First, we used GLIMPSE-1 to identify probable sequence variants in both framework regions (FRs) and CDRs of the heavy and light chains. We combined these variants according to a systematic strategy (**Figure 6A**). We evaluated CDR mutations in pairs, limiting to one chain and one mutation per CDR, and incorporated multiple FR mutations per variant on a single chain at a time. All GLIMPSE-1-generated variants expressed in CHO cells (**Figure 6B**). Affinity testing showed that while many variants had diminished binding to target B, a subset maintained or improved affinity (**Figure 6C**).

In the second stage, we combined all FR mutations with sets of functional CDR mutations across both heavy and light chains. This generated divergent sequences with <90% total identity to the parental clinical antibody. All GLIMPSE-1-generated variants expressed in CHO cells (**Figure 6B**). Many of these extensively modified variants maintained kinetic parameters comparable to the parental antibody’s binding to target B, with several reaching the assay’s limit of quantification for k_off_ alongside the parental molecule (**Figure 6C**). The variants preserved thermostability, contained no additional liabilities compared to the original sequence, and maintained expression yields within expected ranges (**Figure 6B, 6D**).

## 3 Discussion

We developed GLIMPSE-1, a protein language model for practical human antibody engineering. By training solely on human antibody sequences, our model captures the biological patterns and constraints of human antibody repertoires. Since antibodies in the immune system evolve toward optimal heavy-light chain combinations^6^, we trained GLIMPSE-1 on paired chain data. As in somatic hypermutation, GLIMPSE-1 explores a spectrum of mutations, of which many enhance key properties like affinity, yield, and stability, and others do not. This approach aligns with the findings of Shehata *et al*. that human B cell-derived antibodies exhibit distributions of thermal stability, polyreactivity, and hydrophobicity comparable to clinically approved mAbs^10^.

In humanization benchmarking, GLIMPSE-1 performs competitively with Sapiens despite being trained on a dataset much smaller than the training data used in Prihoda *et al*.^8^. This efficiency demonstrates the strategic value of curated, paired human antibody repertoire data over sheer training volume. By focusing exclusively on human paired heavy and light chain variable regions, GLIMPSE-1 avoids the introduction of spurious motifs from non-human and non-antibody proteins. Paired antibody sequences encode biological phenomena that models can harness, such as light chain coherence^6^, to avoid biologically implausible recommendations and minimize polyspecificity^11^. This extends to paratope residues, whereas native antibodies distribute their binding contacts across CDRs while synthetic antibodies rely heavily on CDRH3 contacts^12^.

The benefits of learning from *in vivo* antibody development generalize beyond human sequences, including to llama and mouse with similar improvements in expression, stability, and affinity as in this work^13,14^. This is also true *in vitro*, where cellular quality control in mammalian antibody display filters problematic antibody variants^15^. However, species-specific immunoglobulin adaptations cannot be ignored, such as the CDR-stabilizing DE loop (“CDR4”)^16^. These interactions do not simply arise through somatic hypermutation and affinity maturation. Rather, Shrock *et al*. discovered that species recognize different epitopes due to divergent germline-encoded binding motifs (GRAB motifs), with mice and non-human primates producing different antibody responses than humans to the same antigens^17^. This may explain why “energetic humanization” as performed by Tennenhouse *et al*. appears to outperform traditional humanization approaches^18^.

Human antibody sequences contain evolutionarily optimized features that enhance developability. Foundational work by Zhai *et al*. showed that synthetic libraries designed to mimic natural sequence distributions achieve expression rates of 93% and significantly outperform conventional approaches^19^. While native human antibodies contain chemical liability motifs, those aligned with human germline positions show reduced susceptibility to modification^20^. The majority of therapeutic antibodies occupy a narrow subset of the human antibody developability space, leaving most of the diversity of the human repertoire clinically unutilized^21–23^. Despite this, the benefit of human sequence content is unmistakable and clearly detectable in the framework regions of many approved therapeutic antibodies^24^.

Beyond humanization, we demonstrate the practical utility of GLIMPSE-1 in engineering therapeutic antibodies against multiple targets. Successful engineering requires improvement of one or more attributes without compromising others. Here we show that GLIMPSE-1 generated variants with significant improvements in affinity while maintaining thermostability and expression levels similar to parent sequences. In the case of Target B, these improvements led to sub-nM binding affinities that approached or matched clinical controls, with improvements in both association and dissociation rates. Unsurprisingly, in cases where the paratope is relatively conserved across species, we find that on average improving affinity to the human target protein similarly improves affinity to the non-human protein as well. Our approach is supported by multiple studies that human-trained models can effectively guide engineering and simultaneous optimization of affinity, stability, solubility, and others, albeit with a substantially reduced focus on human antibody data^25–33^. We note in particular work from Frey *et al*., where all successfully engineered variants originated from parental antibodies discovered from *in vivo* immunization campaigns or antibody repertoire sequencing of immunized animals, rather than *de novo* design^34^.

GLIMPSE-1 provides a foundation for engineering therapeutic antibodies using only amino acid sequence data. Somatic hypermutation succeeds *in vivo* despite its inefficiency because billions of B cells can explore sequence space simultaneously at minimal biological cost. Antibody engineering, however, requires time and resources for each variant produced and tested. The framework we employ here utilizes GLIMPSE-1 to enable efficient optimization of antibody humanness and other desirable properties such as binding affinity and developability. Preliminary data from our active learning cycles with GLIMPSE-1 further validates its real-world utility in antibody engineering campaigns, though detailed discussion of these applications falls beyond the scope of this current work.

## 5 Acknowledgments

We sincerely thank the anonymous blood donors whose contributions were essential for this research. Their participation enabled the isolation of B cells critical to Infinimmune’s therapeutic antibody discovery platform and research programs.

## 6 Methods

### 6.1 Training

GLIMPSE-0 is a RoBERTa^5^ encoder-only model, trained against a masked learning modeling (MLM) objective. GLIMPSE-0 was trained and evaluated on a dataset composed of paired Fv sequences from human donors. A number of sequences were withheld from training for evaluation and testing. GLIMPSE-1 employs a modern architecture with custom parameter configurations designed specifically for antibody sequence modeling. GLIMPSE-1 was trained against a masked learning modeling (MLM) objective. GLIMPSE-1 was trained on, evaluated against, and tested against a dataset composed of paired Fv sequences from human donors. Notably donors were disjoint between sets, so as to evaluate and test the model as if it were seeing sequences from a heretofore unobserved donor.

### 6.2 UMAP and FR/CDR Sequence Logos

A 2-dimensional embedding of 633,638 sequence pairs from the proprietary Infinimmune dataset was made by first averaging all final layer activations of GLIMPSE-1 for each of the sequence pairs, then directing the resulting vectors through UMAP^7^ parameterized with 15 neighbors and 2 components. V-gene germline assignments and the total DNA mutations from said germline were inferred for plot annotation using proprietary tooling. Sequence logos^35^ of antibody XY62 (discovered at Infinimmune) were generated by providing the sequence pair to GLIMPSE-1 and transforming the final logits via softmax into likelihoods, then providing the resulting tables to ggseqlogo^36^ for plotting. Germline annotations were provided via alignments given enclone^37^.

### 6.3 Humanization

Twenty clinical antibodies with matched parental mouse sequences from Prihoda *et al*. [8] were identified to benchmark the humanization performance of GLIMPSE-1 against available AbLM variants, including AbLang^38^, AbLang2^39^, BALM-Burbach^4^, ESM-2-ft^4^, BALM-Jing^40^, CurrAb^41^ and Sapiens^8^. Beginning with the parental sequence, and using the output of the previous iteration as input to the next, each model was queried for the single most likely mutation to the input sequence by masking each residue and comparing the softmax-derived log-likelihood of the original residue to what the model predicts under the masked language objective. This mutation, if one existed, was applied to the input sequence, which was subsequently fed into the next iteration. The mutations proposed at each iteration were then compared to the sequence of the clinical molecule, where mutations included in the clinical molecule count toward successful “humanization”. This was done until the model no longer provided likelihood-increasing recommendations or for a maximum of 100 iterations. We tested this approach using three strategies: global, CDR, and paratope. First, we individually masked residues globally across the VH and VL domains of each parental antibody (“global”). Second, we masked FR residues only, defined according to the North definition of CDRs,^42^ allowing each model to evaluate and alter the FR regions but not the CDRs (“CDR”). Finally, for a subset of the antibodies which have solved crystal structures in complex with antigen in the PDB, we masked all non-paratope residues, allowing each model to evaluate and alter any residue that does not contact antigen according to the crystal structure(s) of the clinical antibody and its cognate antigen (“paratope”). The set of paratope-defined antibodies as of April 2025 included certolizumab (5wux), eculizumab (5i5k), ligelizumab (6uqr), omalizumab (5hys, 7shy, 5g64), pembrolizumab (5ggs, 5jxe, 5b8c), solanezumab (4xxd), and tocilizumab (8j6f). Paratope residues for this exercise were defined using ChimeraX version 1.8.^43^ We identified paratope residues using the Contacts workflow to locate residues in the heavy or light chain with ≥ 15.0 Å^2^ buried solvent-accessible surface area in contact with antigen.

### 6.4 Antibody Production

Antibodies were produced at 1 mL scale in ExpiCHO™ cells at Twist Bioscience. Clarified cell culture supernatant containing monoclonal antibodies was purified using Protein A magnetic beads (Pierce™, Thermo Fisher) on the KingFisher™ APEX automated purification system. Briefly, 1 mL of supernatant was incubated with Protein A beads for 1 hr at room temperature with gentle mixing to facilitate antibody binding. Beads were then magnetically harvested and washed twice with phosphate-buffered saline (PBS) to remove unbound proteins and residual media components. Bound antibodies were eluted using 0.2 mM glycine (pH 2.7) and immediately neutralized with 1 M Tris-HCl (pH 8.0). The purified antibodies were stored at 4°C until further use.

Additional antibodies were produced at 1 mL scale in TurboCHO™ at GenScript. Antibodies were either delivered as cell culture supernatants and purified as described above, or purified at GenScript (Protein A purification, eluted in sodium acetate buffer, pH 5.5 with 0.2 M L-Arginine).

### 6.5 Antibody Kinetics Measurements

Surface plasmon resonance (SPR) measurements were taken on a Carterra LSA-XT platform. A PAGHC30M chip was preconditioned with 1 min injections of 25 mM NaOH followed by 10 mM glycine pH 2.0. Antibodies (2 μg/ml) were captured for 10 min. Analytes were prepared at 0.04 nM, 0.2 nM, 1.0 nM, 5.0 nM, and 25 nM. Non-regenerative kinetics were used for each analyte, with 5 minute association and 60 minute dissociation. Regeneration between each series of analyte was performed with 10 mM glycine, pH 2.5, 3 × 45 seconds.

Bio-layer interferometry (BLI) measurements were performed on a Gator Prime Core instrument according to the manufacturer’s recommendations. Briefly, antibodies were diluted in diluent solution (0.1% BSA in PBS) and loaded onto protein A probes without reaching saturation. Following a baselining step, ligand association was measured in 100nM or 200nM antigen solution in diluent. After 120s association, dissociation was measured in diluent solution for an additional 120s. Association rate (k_on_), dissociation rate (k_off_), and dissociation constant (K_D_) were computed using Gator software with Y-axis alignment (Association start, average 10s) and inter-step correction (Association, average 10s) settings turned on.

### 6.6 Antibody Melting Point Measurements

An 8.8 μL aliquot of each sample was loaded into low-volume quartz cuvettes (Unchained Labs) for thermal unfolding analysis using the Uncle instrument (Unchained Labs). The temperature was ramped linearly from 15°C to 95°C at 0.6°C/min, while fluorescence was recorded at excitation wavelengths of 310 nm and 370 nm. Barycentric Mean Fluorescence (BCM) was continuously monitored throughout the temperature ramp. BCM quantifies protein unfolding by measuring the fluorescence-weighted center of mass of the emission spectrum, which shifts as tryptophan and tyrosine residues become solvent-exposed. T_m_ (melting temperature) values were determined from BCM-versus-temperature profiles by calculating the first derivative of the BCM curve, with T_m_ identified as the temperature corresponding to the maximum inflection point. All samples were analyzed in replicates, and T_m_ values are reported as the mean ± standard deviation (SD).

## S Supplemental material

**Table S1:** Machine learning models for antibody engineering.

| Model | Year | Training data | Architecture | Paired | Antibody-specific | Structure required |
| --- | --- | --- | --- | --- | --- | --- |
| Sapiens<sup>8</sup> | 2022 | OAS<sup>2</sup> | RoBERTa<sup>5</sup> | no | yes | no |
| AbLang<sup>38</sup> | 2022 | OAS | RoBERTa | no | yes | no |
| AbLang2<sup>39</sup> | 2024 | OAS | ESM-2 | yes | yes | no |
| ProteinMPNN<sup>44</sup> | 2022 | PDB<sup>1</sup> | Custom | no | no | no |
| ESM-IF1<sup>45</sup> | 2023 | PDB | ESM-IF1 | no | no | yes |
| BALM-Burbach<sup>4</sup> | 2024 | Jaffe *et al.*<sup>6</sup> | RoBERTa | yes | yes | no |
| ESM-2<sup>46</sup> | 2024 | PDB | ESM-2 | no | no | no |
| IgDesign<sup>47</sup> | 2024 | PDB + SAbDab | Custom | no | yes | yes |
| AntiFold<sup>48</sup> | 2024 | OAS + SAbDab | ESM-IF1 | no | yes | yes |
| BALM-Jing<sup>40</sup> | 2024 | OAS | ESM-2 | no | yes | yes |
| CurrAb<sup>41</sup> | 2025 | OAS | ESM-2 | yes | yes | no |

**Figure S1:**
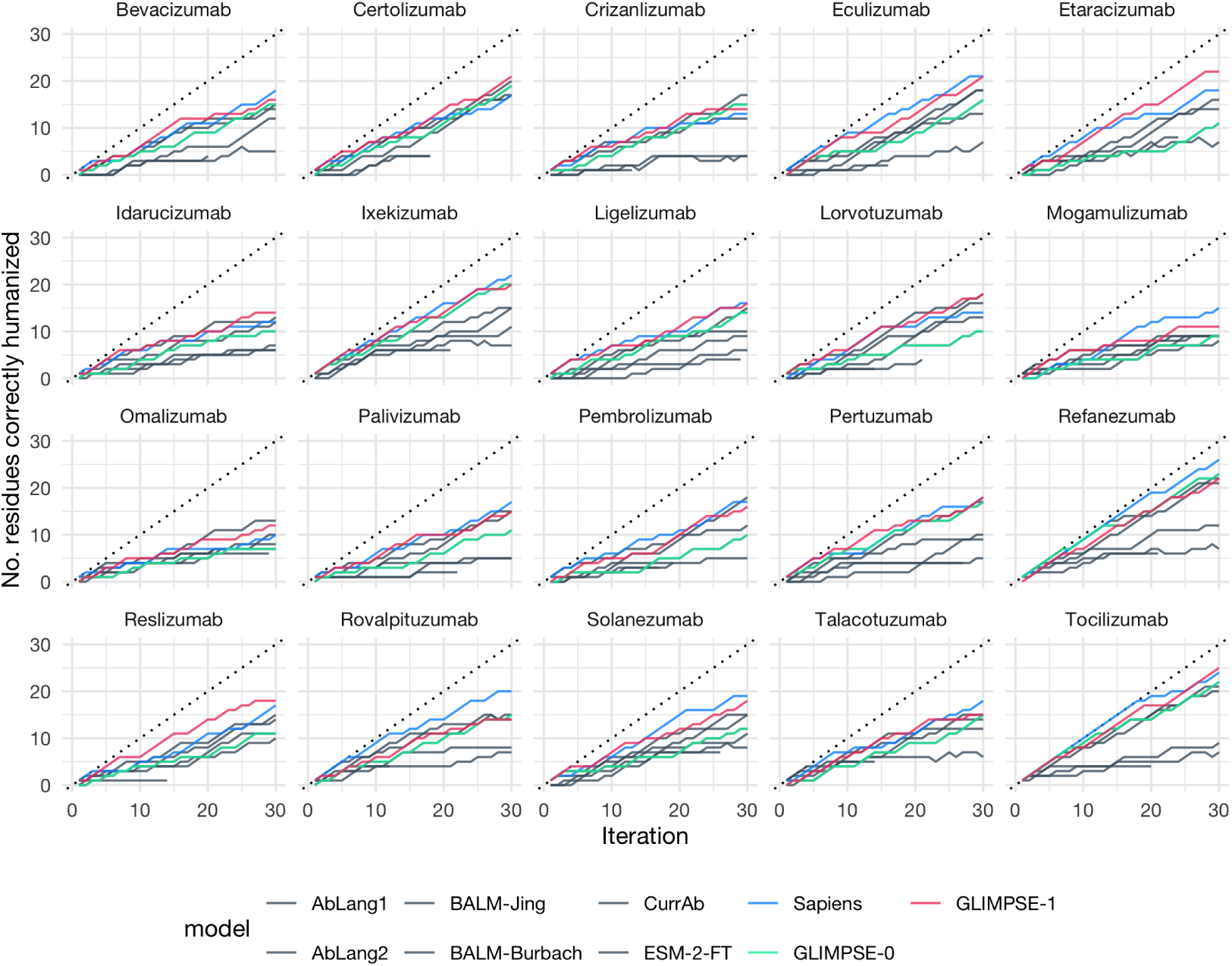
Humanization of 20 mouse-derived clinical stage antibody therapeutics. In contrast to **Figure 3B**, this experiment permitted humanization at all sites.

**Figure S2:**
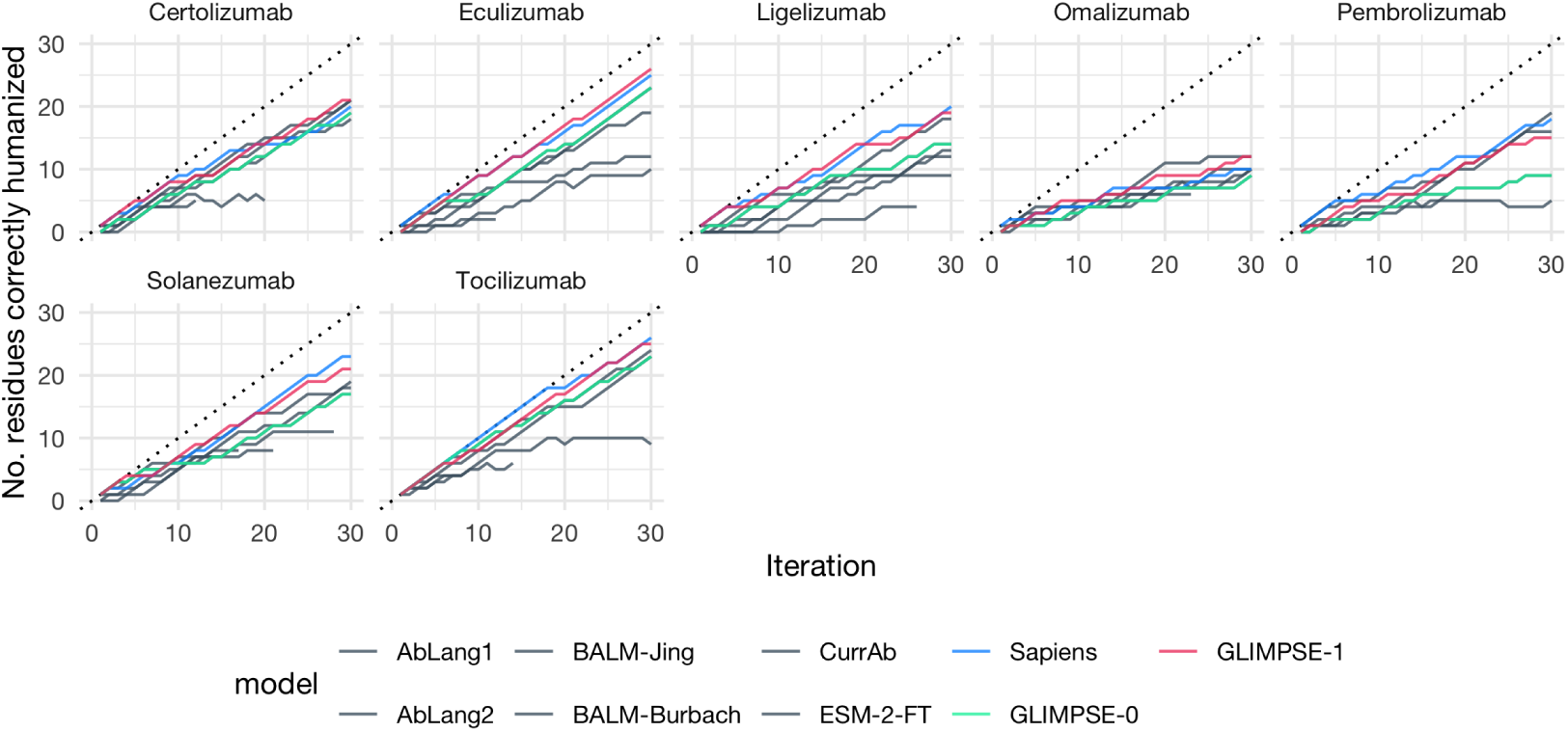
Humanization of 7 mouse-derived clinical stage antibody therapeutics. In contrast to **Figure 3B**, this experiment permitted humanization at any site not found in contact with the target antigen using available structures. See **6.3 Humanization** for details.

